# Differences in polygenic associations with educational attainment between West and East Germany before and after reunification

**DOI:** 10.1101/2024.03.21.585839

**Authors:** D. Fraemke, Y.E. Willems, A. Okbay, G. Wagner, E.M. Tucker-Drob, K.P. Harden, R. Hertwig, P. Koellinger, L. Raffington

## Abstract

Here we examine geographical and historical differences in polygenic associations with educational attainment in East and West Germany around reunification. We test this in n = 1902 25-85-year-olds from the German SOEP-G[ene] cohort. We leverage a DNA-based measure of genetic influence, a polygenic index calculated based on a previous genome-wide association study of educational attainment in individuals living in democratic countries. We find that polygenic associations with educational attainment were significantly stronger among East, but not West, Germans after but not before reunification. Negative control analyses of a polygenic index of height with educational attainment and height indicate that this gene-by-environemt interaction is specific to the educational domain. These findings suggest that the shift from an East German state-socialist to a free-market West German system increased the importance of genetic variants previously identified as important for education.

**One Sentence Summary:** We find that polygenic associations with educational attainment were significantly stronger among East, but not West, Germans after but not before reunification.

## Main Text

After World War II, Germany was divided into two separate states: East Germany became a member of the Soviet-controlled Warsaw Pact and West Germany became a member of NATO, allied with Western democracies. The state-socialist East German regime conducted large-scale institutional reforms aimed at transforming the economic, educational, and other systems to foster opportunities for the working class and reduce intergenerational educational inequality.

Accordingly, East and West German educational systems differed significantly in terms of ideological influence, school types, number of compulsory school years, and student selection (Fischer, 1992; Von Below, 2002). The East German educational system prioritized children of industrial and agricultural workers, provided more academic support to low-achieving students, and less academic support to high-performing students (von Below, 2017). Furthermore, non-meritocratic access restrictions were implemented, which admitted students to higher education based on political affiliation rather than academic performance (Fuchs-Schündeln & Masella, 2016).

In contrast, the West German educational system was (and mostly still is) marked by early school tracking, which combines performance-based measures with input from parents and teachers to sort children into hierarchically organized school tracks at ∼10 years of age. Early school assortment has been found to reproduce intergenerational educational inequality and may account for decreased genetic influence on educational attainment in Germany compared to more liberal educational systems in Norway, Sweden and the US (Baier et al., 2022; Engzell & Raabe, 2023; Lange & von Werder, 2017; Neugebauer, 2010).

With the fall of European state-socialism in 1989/1990, East Germany’s educational ideology of non-meritocratic access restriction was rapidly replaced by West German meritocratic ideology geared towards free-market productivity (Littler, 2017; Rohde, 2023; Solga, 2005). Previous studies have found that the association between education-genetics and educational attainment in Estonia and Hungary increased after the fall of the Soviet Union in Europe (Rimfeld et al., 2018; Ujma et al., 2022). **Here we examine geographical and historical differences in polygenic associations with educational attainment in East and sWest Germany before and after reunification**.

Participants included n = 1902 25-85-year-olds from the German SOEP-G[ene] cohort, of which n = 457 lived in East and n = 1445 in West Germany between 1934 – 2020. We leverage a DNA-based summary measure of genetic influence, a so-called polygenic index of educational attainment. This polygenic index aggregates weighted allele counts that were previously associated with educational attainment in a genome-wide association study of N = 3,037,499 individuals, of which 98.6% completed their education in free-market democracies (EA4; Okbay et al., 2022).

## Results

First, we examined whether the magnitude of the association between education-genetics and educational attainment differed before and after German reunification. We differentiate between individuals who underwent formative school years before and after German reunification by applying an age cut-off of 15 years in 1990 (cf. Rimfeld et al., 2018; see Methods). Educational attainment was quantified in years of education ranging from “compulsory school degree” to “university degree” (range 7 – 18 years). Our education-genetic measure was previously residualized for the top 20 genetic principal components of similarity to ancestral reference groups and genotype batch (Price et al., 2006). We included gender and body mass index (BMI) as covariates in all models, including covariate × environment and covariate × gene interaction terms in the same model that tests gene × environment interaction terms (cf. Keller, 2014).

We find that genetic associations with educational attainment were stronger after German reunification than before in the full sample (see **Model 1** in **Table 1**). This gene-environment interaction remained significant when applying a heteroscedasticity model to assess whether this interaction term is specific to the measured predictor or whether it represents a general pattern of variation in the outcome (**Table 2**; cf. Domingue et al., 2022).

**Table 1.** Model parameter estimates including models with education-genetics, German reunification, and region.

| Term | $\beta$ | SE | $p$ | 95% CI |
| --- | --- | --- | --- | --- |
| **Model 1: Education-genetics x Reunification** |  |  |  |  |
| Education-genetics | 0.26 | 0.03 | < .001*** | [0.20, 0.32] |
| Reunification (post) | 0.44 | 0.07 | < .001*** | [0.30, 0.57] |
| Education-genetics × Reunification (post) | 0.14 | 0.05 | .007** | [0.04, 0.24] |
| **Model 2: Education-genetics x East vs. West Germany** |  |  |  |  |
| Education-genetics | 0.30 | 0.03 | < .001*** | [0.24, 0.37] |
| Region (East Germany) | 0.10 | 0.07 | .143 | [-0.04, 0.24] |
| Education-genetics × Region (East Germany) | 0.07 | 0.05 | .161 | [-0.03, 0.17] |
| **Model 3: Education-genetics x Reunification x East vs. West Germany** |  |  |  |  |
| Education-genetics | 0.26 | 0.03 | < .001*** | [0.19, 0.32] |
| Reunification (Post) | 0.48 | 0.08 | < .001*** | [0.33, 0.63] |
| Region (East Germany) | 0.17 | 0.07 | .024* | [0.02, 0.31] |
| Education-genetics × Reunification (Post) | 0.07 | 0.06 | .241 | [-0.05, 0.20] |
| Education-genetics × Region (East Germany) | 0.01 | 0.06 | .835 | [-0.10, 0.13] |
| Reunification (Post) × Region (East Germany) | -0.06 | 0.12 | .634 | [-0.30, 0.18] |
| Education-genetics × Reunification (Post) × Region (East Germany) | 0.28 | 0.12 | .026* | [0.03, 0.53] |

Note. We controlled for the two covariates gender, body mass index and education-genetics x covariate, reunification x covariate and region x covariate interactions with both covariates (cf. Keller, 2014). Birth year, education-genetics x birth year and region x birth year interactions were included as additional covariates in model 2.

**Table 2.** Heteroscedasticity model parameter estimates.

| | $\tau_1$<br>(SE) | $\pi_0$<br>(SE) | $\pi_1$<br>(SE) | $\lambda_0$<br>(SE) | $\lambda_1$<br>(SE) | $\lambda_2$<br>(SE) | $\xi_E$<br>(SE) | Pr($\xi_E$)<br>(SE) | $\xi_G$<br>(SE) | Pr($\xi_G$)<br>(SE) |
| --- | --- | --- | --- | --- | --- | --- | --- | --- | --- | --- |
| Education-genetics<br>x Reunification | 0.26<br>(0.05) | 0.28<br>(0.05) | 0.16<br>(0.06) | 0.93<br>(0.02) | 0.06<br>(0.04) | 0.09<br>(0.09) | -0.13<br>(0.11) | .015* | -0.12<br>(0.09) | .013* |
| Education-genetics<br>x Birth Year | 0.20<br>(0.03) | 0.34<br>(0.02) | 0.10<br>(0.01) | 0.95<br>(0.02) | 0.04<br>(0.02) | 0.09<br>(0.02) | -0.08<br>(0.06) | .006** | -0.08<br>(0.05) | .005** |

Note. Parameter estimates obtained from an environmental heteroscedasticity model are $\tau_1$ , the main effect of the measured environment, $\pi_0$ , the main effect of education-genetics, $\pi_1$ , the gene-by-environment interaction effect, $\lambda_0$ , the main effect of error term, and $\lambda_1$ , the interaction between the environment and error term, indexing environmental heteroscedasticity. The parameter estimate obtained from a genetic heteroscedasticity model is $\lambda_2$ , the interaction between education-genetics and the error term indexing genetic heteroscedasticity. $\xi_E$ and $\xi_G$ are test statistics derived from the corresponding model following the formula $\xi_E \equiv \pi_0 \lambda_1 - \pi_1 \lambda_0$ and $\xi_G \equiv \pi_0 \lambda_2 - \pi_1 \lambda_0$ . Significant $\chi^2$ -tests of $\xi$ indicate that the gene-by-environment interaction is not driven by heteroscedasticity in the environment or education-genetics (cf. Domingue et al., 2022). H0: $\xi = 0$ ; gene-by-environment interaction is driven by dispersion in outcome related to G or E. H1: $\xi \neq 0$ ; GxE interaction is not driven by dispersion in outcome related to G or E.

Second, we examined whether the strength of genetic associations with educational attainment differed between East and West Germany in the full sample. They did not (see **Model 2** in **Table 2**). There were also no statistically significant mean differences in the education PGI (*t*(838.47) = 0.80, *p* = 0.42) or educational attainment (*t*(828.1) = -1.03, *p* = 0.30) between East and West Germans (see **Supplemental Figure S1**).

Third, we examined whether the historical period differences in genetic associations with educational attainment (before versus after German reunification) differed in East versus West Germany. We find that the increase in education-genetic association with educational attainment after German reunification significantly differed by region (*i*.*e*., East vs West; see **Model 3** in **Table 1**). Post-hoc analyses that split the sample by region suggest that this interaction was driven by a post-reunification increase in the magnitude of the association between education-genetics and educational attainment in East Germany (reunification × education-genetics: β = 0.35 [95% CI: 0.15, 0.54], *p* < 0.001), not West Germany (reunification × education-genetics: β = 0.07 [95% CI: -0.06, 0.2], *p* = 0.25).

**Figure 1 Panel A** plots the incremental R^2^ values of the association between education-genetics and educational attainment by region and reunification. While effect size estimates were similar between East and West Germany before reunification and remained stable in West Germany (R^2^_pre_ = 7.4% [95% CI: 5, 11%] to R^2^_post_ = 8.3% [3, 15%]), there was a substantial post-reunification increase in the magnitude of the education-genetics association in East Germany (R^2^_pre_ = 9.1% [4, 15%] to R^2^_post_ = 37.6% [24, 50%]). In comparison, the SNP-heritability estimates for years of education in Estonia were 18% (95% CI: 12, 24%) before the fall of the Soviet Union and increased to 37% (95% CI: 10, 64%) after. We caution that small sample sizes, such as our post-reunification subsample, tend to overestimate effect size estimates (Hedges & Olkin, 1985). See Supplemental **Figure S2** for scatterplots.

**Figure 1.**
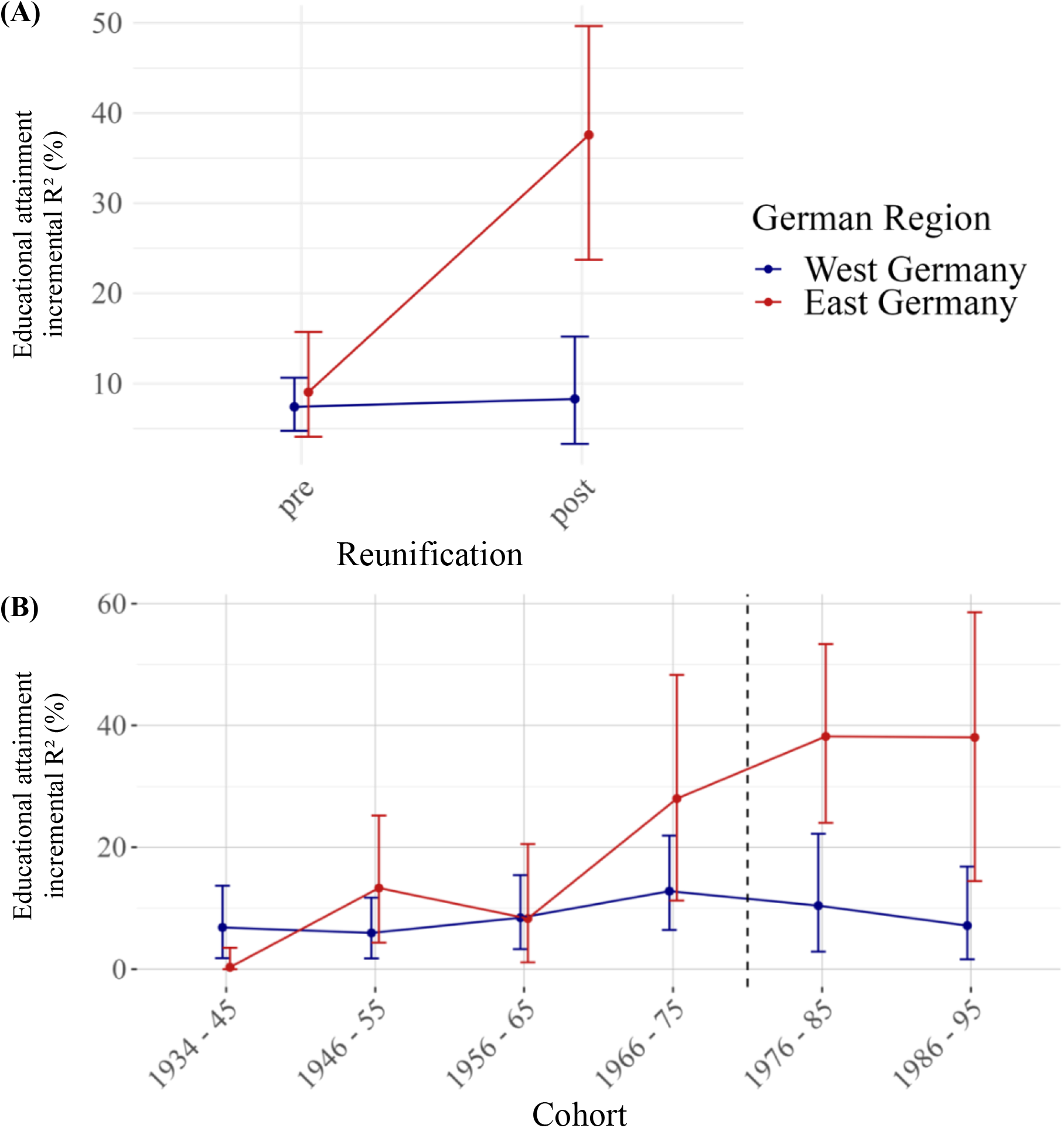
Effect size estimates of the association between education-genetics and educational attainment by region. **Panel A** plots the model-implied incremental R^2^ of education-genetics (*i*.*e*., EA4) and educational attainment before and after German reunification in East and West Germany. **Panel B** plots the model-implied incremental R^2^ of education-genetics and educational attainment by birth year in East and West Germany. We caution that small sample sizes, such as our post-reunification subsample, tend to overestimate effect size estimates (Hedges & Olkin, 1985). Error bars indicate 95% bootstrapped confidence intervals with 2,000 replications. The vertical dashed line divides cohorts which were older (left of the line) and younger (right of the line) than 15 years at the time of German reunification in 1990.

Since German reunification denotes a single timepoint historical event, it can be considered a historically bound categorical version of birth year. Thus, we examined whether the above gene-environment interaction replicated using a birth year variable rather than German reunification. In line with our above results, the magnitude of the association between education-genetics and educational attainment increased in younger Germans (**Model 1b** in **Table 3**). This gene-environment interaction also remained significant when applying a heteroscedasticity model (**Table 2**). We find that the increase in genetic association with educational attainment in younger Germans also differed by region (**Model 3b** in **Table 3**). Post-hoc analyses that split the sample by region suggest that this interaction was driven by an increase in genetic association in younger East Germans (birth year × education-genetics: β = 0.24 [0.13, 0.35], *p* < 0.001), not younger West Germans (birth year × education-genetics: β = 0.07 [-0.01, 0.14], *p* = 0.07).

**Table 3.** Model parameter estimates including models with education-genetics, birth year, and region.

| Term | $\beta$ | SE | p | 95% CI |
| --- | --- | --- | --- | --- |
| **Model 1b:** Education-genetics x Birth Year |  |  |  |  |
| Education-genetics | 0.32 | 0.03 | < .001*** | [0.27, 0.38] |
| Birth Year | 0.34 | 0.04 | < .001*** | [0.27, 0.42] |
| Education-genetics × Birth Year | 0.10 | 0.03 | .001** | [0.04, 0.16] |
| **Model 3b:** Education-genetics x Birth Year x East vs. West Germany |  |  |  |  |
| Education-genetics | 0.30 | 0.03 | < .001*** | [0.23, 0.36] |
| Birth Year | 0.38 | 0.04 | < .001*** | [0.30, 0.47] |
| Region (East Germany) | 0.14 | 0.07 | .057 | [-0.00, 0.28] |
| Education-genetics × Birth Year | 0.07 | 0.04 | .058 | [-0.00, 0.14] |
| Education-genetics × Region (East Germany) | 0.12 | 0.06 | .027* | [0.01, 0.23] |
| Birth Year × Region | -0.11 | 0.07 | .132 | [-0.25, 0.03] |
| Education-genetics × Birth Year × Region (East Germany) | 0.17 | 0.07 | .016* | [0.03, 0.31] |

*Note.* We controlled for the two covariates sex, body mass index, and education-genetics × covariate, birth year × covariate and region × covariate interactions with both covariates.

**Figure 1 Panel B** plots the incremental R^2^ values of the association between education-genetics and educational attainment by region and birth year bins. Models including both German reunification and birth year interactions in the same model indicated that their effects are too collinear to distinguish (see **Supplemental Results**). Lastly, negative control analyses of polygenic index of height with educational attainment and height indicate that these gene by reunification interactions are specific to the educational domain (see **Supplemental Results**).

## Discussion

We examined geographical and historical differences in the association between a DNA-based measure of education-genetics and educational attainment in East and West Germany before and after reunification in n = 1902 25-85-year-olds from the SOEP-G cohort. We found that after German reunification the magnitude of the association between education-genetics and educational attainment increased in East Germans, who experienced a monumental shift from a state-socialist to a free-market West German system. This aligns with previous reports that the association between education-genetics and educational attainment increased in Estonia and Hungary after the fall of the Soviet Union in Europe (Rimfeld et al., 2018; Ujma et al., 2022). Notably, our time-concurrent regional contrast between East and West Germany substantially reduces the likelihood of cohort-related confounds that could affect genetic associations. Moreover, negative control analyses of a polygenic index of height with educational attainment and self-reported height indicate that this G x E interaction is specific to the educational domain. Collectively, these findings are consistent with the interpretation that there was an amplification of education-genetic associations post-Soviet Union/German reunification rather than a Soviet-era suppression of genetic effects relative to West Germany.

Our results indicate that education-genetic associations changed with the emergence of novel social norms and structures. Following the collapse of socialist elites, the post-reunification generation of East Germans found themselves uniquely liberated from the constraints of ideological selection and parental social status (Klein et al., 2019). As old social norms begin to be replaced by new norms, individuals have greater choice in their outcomes: they can continue with the status quo, or they can adopt the newer social tides (Briley et al 2015). Because educational behaviors are affected by genetically influenced dispositions, genetic influences are predicted to increase during periods of social change marked by increasing social, educational, and economic opportunities (Engzell & Tropf, 2019; Raffington et al., 2020; Tucker-Drob et al., 2013).

We caution that polygenic indices of educational attainment do not capture *all* education-genetic effects and reflect a mixture of direct genetic influence (*e*.*g*., one’s disposition to academic persistence), indirect genetic influence (*e*.*g*., parental nurture effects on child education), but also socially-stratified environmental differences between families (*e*.*g*., dynastic social processes; Aikins et al., 2024; Malanchini et al., 2023; Nivard et al., 2024; Wertz et al., 2023). It is possible that genetic propensity for traits such as social conformity were more relevant to educational attainment in pre-unification East Germany, and these factors are not captured by this polygenic index of educational attainment, which was primarily based on genetic discovery in people living in free-market Western democracies. Yet, our finding that genetic associations were similar between East and West Germany before reunification is more consistent with the interpretation that genetic influences became more important during this social transition than the interpretation that different genes mattered in East versus West Germany. Future studies could corroborate this interpretation by probing whether educational performance and aspirations were more predictive of educational attainment in East compared to West Germany shortly after reunification.

A meritocratic-oriented educational system may have both desirable and unwanted effects on social justice and cohesion. It is generally considered desirable for a society to have individuals who excel in roles such as caregivers, public servants, and pilots. Thus, matching genetically-influenced skills and preferences to corresponding educational training and vocations is beneficial for society-at-large and may reflect a more equal society that does not restrict access to education on the basis of class, gender, or similar (Raffington et al., 2020). Yet, meritocratic selection that leads to substantially higher monetary and health rewards on the basis of genetically-influenced performance and aspirations could have unintended consequences that amplify intergenerational social inequality and threaten social cohesion (Bloodworth, 2016; Harden, 2021; Knigge et al., 2022; Rimfeld et al., 2018). Tackling the downstream individual and intergenerational inequities in income, health, and life opportunities that arise through differences in educational attainment is thus of utmost importance to foster social justice and coherence in meritocratic-oriented societies.

## Methods

### Participants

The Socio-Economic Panel (SOEP) is a population-based, multi-generational survey study (Goebel et al., 2019). 6,576 SOEP participants were randomly selected and invited to participate in buccal DNA genotyping as part of the SOEP-Gene subsample (SOEP-G; Koellinger et al., 2023). In total, education-genetics are available for n = 2,262 adults (M_age_ = 56.13, SD_age_ = 18.72, 54% female), with 98% of participants showing high genetic similarity to European reference groups (see Koellinger et al., 2023). Education-genetic analyses were restricted to participants with high genetic similarity to European reference groups, which formed the basis of the GWAS discovery sample, to avoid the risk of confounding due to population stratification (Price et al., 2006). See **Supplemental Methods** for DNA preprocessing.

Present analyses included n = 1902 25-85-year-olds from the German SOEP-G[ene] cohort, of which n = 457 lived in East Germany and n = 1445 in West Germany between the years 1949 – 2020. We differentiate between individuals who underwent formative school years before and after German reunification by applying an age cut-off of 15 years in 1990 (cf. Rimfeld et al., 2018). Among East Germans this cut-off resulted in n = 350 individuals that turned 15 before the reunification and n = 107 after. Among West Germans n = 1122 turned 15 before reunification and n = 323 after. Individuals born before 1934 were excluded (n = 86), as they turned 15 before Germany was officially separated in 1949. **Table 4** reports descriptive statistics and **Table 5** main variables of interest.

**Table 4.** Descriptive statistics of variables in East and West Germans.

| Variable | West |  |  | East |  |  |
| --- | --- | --- | --- | --- | --- | --- |
|  | Mean | SD | [min; max] | Mean | SD | [min; max] |
| Female (in %) | 53.96 |  |  | 53.89 |  |  |
| Age | 57.67 | 16.26 | [25; 85] | 58.41 | 15.75 | [25.2; 84.8] |
| BMI | 26.99 | 5.36 | [14.6; 64.9] | 27.54 | 5.71 | [17.3; 57.1] |
| Education-genetics | 0.01 | 0.98 | [-3.9; 3.5] | -0.05 | 0.98 | [-3; 3.3] |
| Years of Education | 12.52 | 2.76 | [7; 18] | 12.66 | 2.54 | [7; 18] |
| strict QC pass (in %) | 91.08 |  |  | 92.21 |  |  |

*Note.* QC = Quality Control of genetic data. Education-genetic index was z-standardized.

**Table 5.** Main variables of interest.

| Construct | Description |
| --- | --- |
| Educational Attainment | Educational Attainment was assessed in number of years of education an individual obtained on a scale ranging from “compulsory school degree” (min = 7 years) to “university degree” (max = 18 years). Tertiary education in years was added to the years spent in secondary education (e.g., +1.5 years for vocational training or +5 years for a university degree; Goebel et al. 2019). |
| East vs West Germany | Whether an individual lived in East or West Germany during or after the reunification was assessed by two items in the SOEP dataset. The individuals indicated, whether they lived in East or West Germany before the fall of the Berlin Wall in 1989 (ppfad/loc1989). For individuals who were born after or had missing data on this variable, their household location was used (hbrutto/sampreg; Goebel et al., 2019). |
| Education-genetics | The polygenic index of educational attainment was computed based on the most recent genome-wide association study of educational attainment in a sample of ~ 3 million individuals of high genetic similarity to European reference groups (EA4; Okbay et al., 2022). EA4 was computed using SBayesR using the default settings (Lloyd-Jones et al., 2019). EA4 was residualized for the top 20 genetic principal components of ancestry and genotype batch (Price et al., 2006). |
| Covariates | Age, gender<br>Body Mass Index (BMI) was computed by transforming self-reported height (in cm) and self-reported weight (in kg) to sex- and age-normed z-scores. |

## Supporting information

Supplemental Results

## Funding Acknowledgements

During his work on this paper, DF was a pre-doctoral fellow of the International Max Planck Research School on the Life Course (LIFE, www.imprs-life.mpg.de; participating institutions: Max Planck Institute for Human Development, Freie Universität Berlin, Humboldt-Universität zu Berlin, University of Michigan, University of Virginia, University of Zurich). EMTD was supported by the National Institutes of Health (NIH) grants RF1AG073593, R01MH120219, and R01HD092548. EMTD and KPH are Faculty Research Associate of the Population Research Center (PRC), which is supported by a NIH grant P2CHD042849. EMTD is Faculty Research Associate of the Center for Aging and Populations Sciences (CAPS), which is supported by NIH grant P30AG066614. KPH was supported by NIH grants R01HD083613 and R01HD092548. PK received funding from the University of Amsterdam. RH and GGW received funding from the German Research Foundation DFG and the Max Planck Society. LR is faculty member at LIFE and received funding from the Max Planck Society. Funders/support had no role in the design and conduct of the study; collection, management, analysis, and interpretation of the data; preparation, review, or approval of the manuscript; and decision to submit the manuscript for publication.

