## Supplemental Methods for "Differences in polygenic associations with educational attainment between West and East Germany before and after reunification"

**Genetic data**

Buccal swabs and Isohelix IS SK-1S Dri-Capsules were used to collect DNA data. DNA extraction and methylation profiling were conducted at the Erasmus Medical Center in the Netherlands by the Human Genomics Facility (HuGe-F). The use of the DNA samples received ethical approval by The Vrije Universiteit Amsterdam, School of Business and Economics (application number 20181018.1.pkr730) and the Max Planck Society (application number 2019_16). Genotyping was conducted using the Illumina Infinium Global Screening Array-24 v3.0 BeadChips. Genotypes were subject to quality control excluding participants with sex mismatch, with per-chromosome missingness of more than 50%, and with excess heterozygosity/homozygosity.

The Haplotype Reference Consortium reference panel (r.1.1) for imputation was used with imputation accuracy (R^2^) greater than 0.1. Approximately 66% of the imputed SNPs were rare with minor allele frequencies (MAF) smaller than 0.01 and ~24% SNPs were common. The average imputation accuracy in the data was 0.66, with higher imputation accuracy for common SNPs (MAF>0.05) with an average imputation accuracy of 0.92. To control for population stratification, the first 20 principal components (PCs) were computed for individuals with high genetic similarity to European reference groups, based on ~160,000 approximately independent SNPs with imputation accuracy ≥70% and MAF≥0.01 (see Koellinger et al., 2023).
