## Supplemental Results for "Differences in polygenic associations with educational attainment between West and East Germany before and after reunification"

In a model that included both German reunification and birth year interactions with education-genetics, only the gene x birth year interaction remained significant (**Model 1 in Table S1**). In a model that included both German reunification x education-genetics x East/West and birth year x education-genetics x East/West interactions, neither of the interactions was statistically significant (**Model 3 in Table S1**). This indicates that the effects of German reunification and birth year are too collinear to distinguish.

We conducted several sensitivity analyses to probe the robustness of our main results. We examined if genetic relatedness within families or genotyping quality introduced unwanted bias in the results. Excluding individuals who had either siblings or parents in the sample (n = 17) did not affect results (**Table S2**). Excluding individuals that had not passed a strict genotyping quality control (QC) pipeline (n = 172; Koellinger et al., 2023) did not substantially affect model parameters (**Table S3)**. Moreover, inclusion of BMI as a covariate did not substantially affect the results (**Table S4**).

Lastly, we conducted negative control analyses of polygenic index of height (Wood et al., 2014) with educational attainment and height to test for a general pattern of elevated genetic influence. There were no associations of height-genetics with educational attainment (**Model 3 in Table S5**). Moreover, associations of height-genetics with self‑reported height did not differ pre-/post German reunification, between East and West Germany, or birth year (**Model 3 in Table S6; Figure S3**). Hence, gene by reunification/region interactions were specific to the educational domain.

| **Table S1.**  Full model results with both German reunification and birth year in one model. | | | | |
| --- | --- | --- | --- | --- |
| **Term** | **β** | **SE** | ***p*** | **95% CI** |
| **Model 1:**  Education-genetics x Birth Year and  Education-genetics x Reunification | | | | |
| Education-genetics | 0.33 | 0.04 | < .001*** | [0.25, 0.40] |
| Birth Year | 0.37 | 0.06 | < .001*** | [0.24, 0.49] |
| Reunification (Post) | -0.05 | 0.11 | .615 | [-0.26, 0.16] |
| Education-genetics ×  Birth Year | 0.10 | 0.05 | .026* | [0.01, 0.20] |
| Education-genetics ×  Reunification (Post) | -0.01 | 0.08 | .923 | [-0.17, 0.15] |
| **Model 3:**  Education-genetics x Birth Year x East vs. West Germany and  Education-genetics x Reunification x East vs. West Germany | | | | |
| Education-genetics | 0.32 | 0.04 | < .001*** | [0.23, 0.41] |
| Region (East Germany) | 0.05 | 0.10 | .611 | [-0.14, 0.24] |
| Education-genetics ×  Region (East Germany) | 0.08 | 0.09 | .373 | [-0.09, 0.25] |
| Reunification Terms |  |  |  |  |
| Reunification (Post) | -0.14 | 0.12 | .269 | [-0.38, 0.10] |
| Reunification (Post) ×  Education-genetics | -0.08 | 0.10 | .408 | [-0.27, 0.11] |
| Reunification (Post) ×  Region (East Germany) | 0.25 | 0.20 | .207 | [-0.14, 0.64] |
| Reunification (Post) ×  Education-genetics ×  Region (East Germany) | 0.14 | 0.20 | .487 | [-0.25, 0.53] |
| Birth Year Terms |  |  |  |  |
| Birth Year | 0.44 | 0.07 | < .001*** | [0.30, 0.58] |
| Birth Year ×  Education-genetics | 0.10 | 0.06 | .071 | [-0.01, 0.21] |
| Birth Year ×  Region (East Germany) | -0.22 | 0.12 | .057 | [-0.45, 0.01] |
| Birth Year ×  Education-genetics ×  Region (East Germany) | 0.12 | 0.12 | .313 | [-0.11, 0.34] |
| *Note.* We controlled for the two covariates sex, BMI and education-genetics × covariate, birth year × covariate, reunification × covariate and region × covariate interactions with both covariates. | | | | |

| **Table S2**  Full model results in unrelated individuals. | | | | |
| --- | --- | --- | --- | --- |
| **Term** | **β** | **SE** | ***p*** | **95% CI** |
| **Model 1:** Education-genetics x Reunification | | | | |
| Education-genetics | 0.26 | 0.03 | < .001*** | [0.20, 0.32] |
| Reunification (post) | 0.44 | 0.07 | < .001*** | [0.31, 0.58] |
| Education-genetics ×  Reunification (post) | 0.14 | 0.05 | .007** | [0.04, 0.25] |
| **Model 2**: Education-genetics x East vs. West Germany | | | | |
| Education-genetics | 0.30 | 0.03 | < .001*** | [0.24, 0.37] |
| Region (East Germany) | 0.10 | 0.07 | .143 | [-0.04, 0.24] |
| PGI-Education ×  Country (East Germany) | 0.07 | 0.05 | .161 | [-0.03, 0.17] |
| **Model 3:** PGI-Education x Reunification x East vs. West Germany | | | | |
| PGI-Education | 0.26 | 0.03 | < .001*** | [0.19, 0.32] |
| Reunification (Post) | 0.50 | 0.08 | < .001*** | [0.35, 0.65] |
| Region (East Germany) | 0.17 | 0.07 | .023* | [0.02, 0.31] |
| PGI-Education × Reunification (Post) | 0.09 | 0.06 | .169 | [-0.04, 0.22] |
| PGI-Education ×  Region (East Germany) | 0.01 | 0.06 | .831 | [-0.10, 0.13] |
| Reunification (Post) × Region (East Germany) | -0.08 | 0.12 | .541 | [-0.32, 0.17] |
| PGI-Education × Reunification (Post) × Region (East Germany) | 0.27 | 0.13 | .033* | [0.02, 0.51] |
| *Note.* Model parameter after excluding individuals with samples that did not pass strict quality control of genetic data. We controlled for the two covariates gender, BMI and education-genetics x covariate, reunification x covariate and region x covariate interactions with both covariates (cf. Keller, 2014). | | | | |

| **Table S3**  Full model results in subsample passing strict DNA quality control. | | | | |
| --- | --- | --- | --- | --- |
| **Term** | **β** | **SE** | ***p*** | **95% CI** |
| **Model 1:** Education-genetics x Reunification | | | | |
| Education-genetics | 0.27 | 0.03 | < .001*** | [0.20, 0.33] |
| Reunification (post) | 0.24 | 0.05 | < .001*** | [0.15, 0.34] |
| Education-genetics ×  Reunification (post) | 0.13 | 0.06 | .018* | [0.02, 0.24] |
| **Model 2**: Education-genetics x East vs. West Germany | | | | |
| Education-genetics | 0.31 | 0.04 | < .001*** | [0.24, 0.38] |
| Region (East Germany) | 0.12 | 0.07 | .102 | [-0.02, 0.26] |
| Education-genetics ×  Country (East Germany) | 0.06 | 0.05 | .290 | [-0.05, 0.16] |
| **Model 3:** Education-genetics x Reunification x East vs. West Germany | | | | |
| Education-genetics | 0.27 | 0.04 | < .001*** | [0.20, 0.34] |
| Reunification (Post) | 0.49 | 0.08 | < .001*** | [0.34, 0.65] |
| Region (East Germany) | 0.19 | 0.08 | .014* | [0.04, 0.33] |
| Education-genetics × Reunification (Post) | 0.07 | 0.07 | .304 | [-0.06, 0.20] |
| Education-genetics ×  Region (East Germany) | -0.01 | 0.06 | .882 | [-0.13, 0.11] |
| Reunification (Post) × Region (East Germany) | -0.07 | 0.13 | .555 | [-0.32, 0.17] |
| Education-genetics × Reunification (Post) × Region (East Germany) | 0.28 | 0.13 | .031* | [0.03, 0.53] |
| *Note.* Model parameter after excluding individuals with samples that did not pass strict quality control of genetic data. We controlled for the two covariates gender, BMI and education-genetics x covariate, reunification x covariate and region x covariate interactions with both covariates (cf. Keller, 2014). | | | | |

| **Table S4**  Full model results without covariate control for body mass index. | | | | |
| --- | --- | --- | --- | --- |
| **Term** | **β** | **SE** | ***p*** | **95% CI** |
| **Model 1:** Education-genetics x Reunification | | | | |
| Education-genetics | 0.28 | 0.03 | < .001*** | [0.22, 0.34] |
| Reunification (post) | 0.46 | 0.07 | < .001*** | [0.33, 0.60] |
| Education-genetics ×  Reunification (post) | 0.15 | 0.05 | .003** | [0.05, 0.26] |
| **Model 2**: Education-genetics x East vs. West Germany | | | | |
| Education-genetics | 0.31 | 0.04 | < .001*** | [0.24, 0.38] |
| Region (East Germany) | 0.12 | 0.07 | .102 | [-0.02, 0.26] |
| Education-genetics ×  Country (East Germany) | 0.06 | 0.05 | .290 | [-0.05, 0.16] |
| **Model 3:** Education-genetics x Reunification x East vs. West Germany | | | | |
| Education-genetics | 0.28 | 0.03 | < .001*** | [0.22, 0.35] |
| Reunification (Post) | 0.14 | 0.07 | .051 | [-0.00, 0.28] |
| Region (East Germany) | 0.50 | 0.08 | < .001*** | [0.35, 0.65] |
| Education-genetics × Reunification (Post) | -0.02 | 0.06 | .777 | [-0.13, 0.10] |
| Education-genetics ×  Region (East Germany) | 0.10 | 0.06 | .117 | [-0.02, 0.22] |
| Reunification (Post) × Region (East Germany) | -0.04 | 0.12 | .756 | [-0.28, 0.20] |
| Education-genetics × Reunification (Post) × Region (East Germany) | 0.27 | 0.12 | .031* | [0.02, 0.52] |
| *Note.* We controlled for the covariate gender, education-genetics x gender, reunification x gender and Region x gender (cf. Keller, 2014). | | | | |

|  | **Table S5**  Full model results of height-genetics on educational attainment. | | | | |
| --- | --- | --- | --- | --- | --- |
| **Term** | | **β** | **SE** | ***p*** | **95% CI** |
| **Model 3:**  Height-Genetics x Birth Year x East vs. West Germany and  Height-Genetics x Reunification x East vs. West Germany | | | | | |
| Height-Genetics | | -0.06 | 0.05 | .229 | [-0.15, 0.04] |
| Region (East Germany) | | -0.01 | 0.11 | .916 | [-0.22, 0.20] |
| Height-Genetics ×  Region (East Germany) | | 0.07 | 0.10 | .497 | [-0.12, 0.26] |
| Reunification Terms | |  |  |  |  |
| Reunification (Post) | | 0.05 | 0.11 | .686 | [-0.18, 0.27] |
| Reunification (Post) ×  Height-Genetics | | 0.10 | 0.10 | .317 | [-0.10, 0.31] |
| Reunification (Post) ×  Region (East Germany) | | 0.15 | 0.21 | .483 | [-0.27, 0.57] |
| Reunification (Post) ×  Height-Genetics ×  Region (East Germany) | | -0.04 | 0.21 | .836 | [-0.46, 0.37] |
| Birth Year Terms | |  |  |  |  |
| Birth Year | | 0.27 | 0.06 | < .001*** | [0.15, 0.38] |
| Birth Year ×  Height-Genetics | | -0.10 | 0.06 | .074 | [-0.22, 0.01] |
| Birth Year ×  Region (East Germany) | | -0.25 | 0.13 | .046* | [-0.50, -0.00] |
| Birth Year ×  Height-Genetics ×  Region (East Germany) | | 0.03 | 0.13 | .805 | [-0.22, 0.28] |
| *Note.* We controlled for the covariate gender, education-genetics × gender, birth year × gender, reunification × gender and region ×gender interactions. | | | | | |

| **Table S6**  Full model results of associations between height-genetics and height. | | | | |
| --- | --- | --- | --- | --- |
| **Term** | **β** | **SE** | **p** | **95% CI** |
| **Model 3c:**  Height-Genetics x Birth Year x East vs. West Germany and  Height-Genetics x Reunification x East vs. West Germany | | | | |
| Height-Genetics | 0.33 | 0.03 | < .001*** | [0.27, 0.39] |
| Region (East Germany) | -0.21 | 0.07 | .001** | [-0.34, -0.09] |
| Height-Genetics ×  Region (East Germany) | -0.03 | 0.06 | .632 | [-0.15, 0.09] |
| Reunification Terms |  |  |  |  |
| Reunification (Post) | -0.29 | 0.07 | < .001*** | [-0.43, -0.16] |
| Reunification (Post) ×  Height-Genetics | -0.04 | 0.06 | .515 | [-0.17, 0.08] |
| Reunification (Post) ×  Region (East Germany) | 0.20 | 0.13 | .131 | [-0.06, 0.45] |
| Reunification (Post) ×  Height-Genetics ×  Region (East Germany) | 0.00 | 0.13 | .989 | [-0.25, 0.26] |
| Birth Year Terms |  |  |  |  |
| Birth Year | 0.41 | 0.04 | < .001*** | [0.34, 0.48] |
| Birth Year ×  Height-Genetics | 0.03 | 0.04 | .359 | [-0.04, 0.10] |
| Birth Year ×  Region (East Germany) | 0.03 | 0.08 | .680 | [-0.12, 0.18] |
| Birth Year ×  Height-Genetics ×  Region (East Germany) | -0.03 | 0.08 | .699 | [-0.18, 0.12] |
| *Note.* We controlled for the covariate gender, education-genetics × gender, birth year × gender, reunification × gender and region ×gender interactions. | | | | |

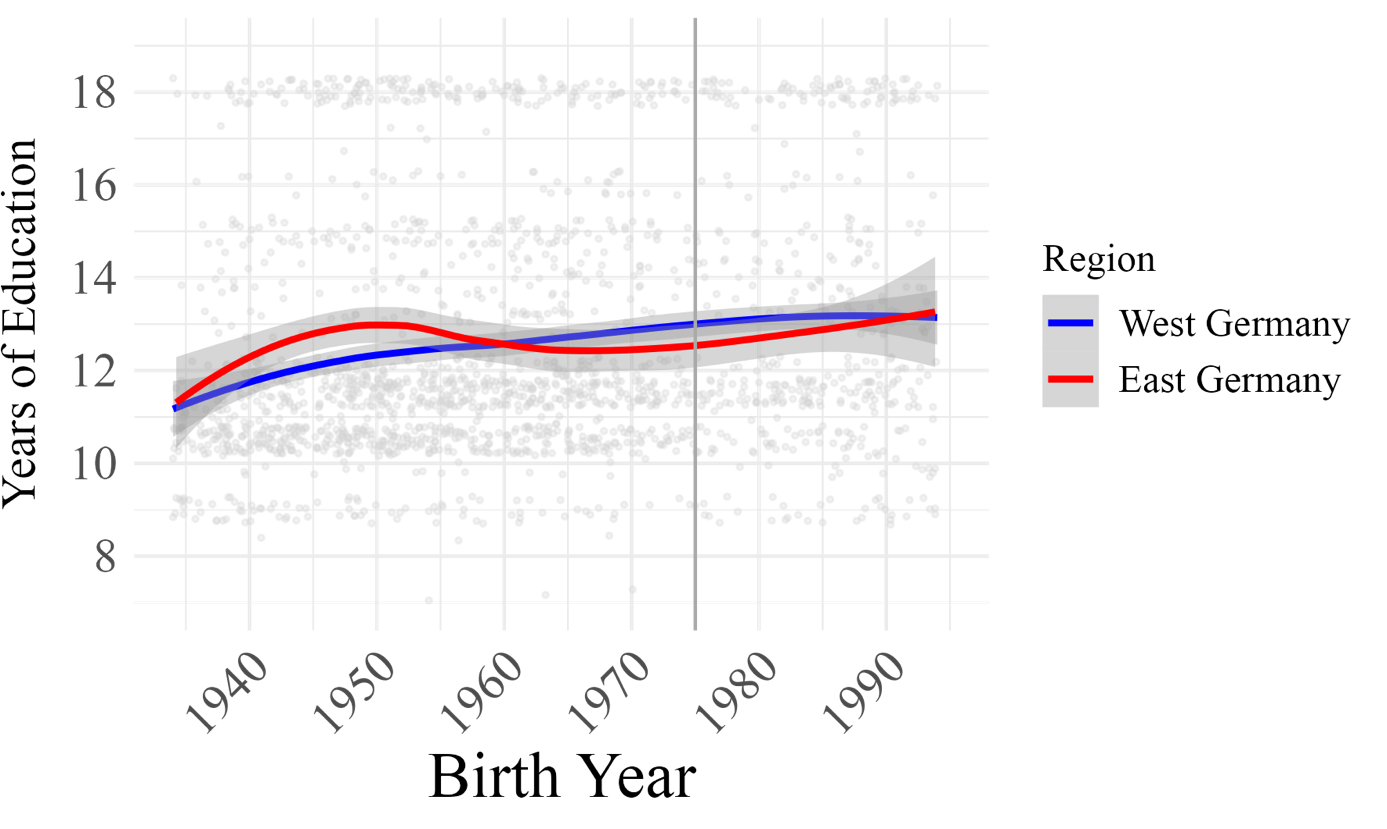

**Figure S1.** **Years of educational attainment by birth year in East and West Germany.** The first vertical line separates individuals who turned 15 years before and after German reunification in 1990. Clustering random noise is added to years of education in this figure for visualization purposes.

East Germany

West Germany

Before German reunification

After German reunification

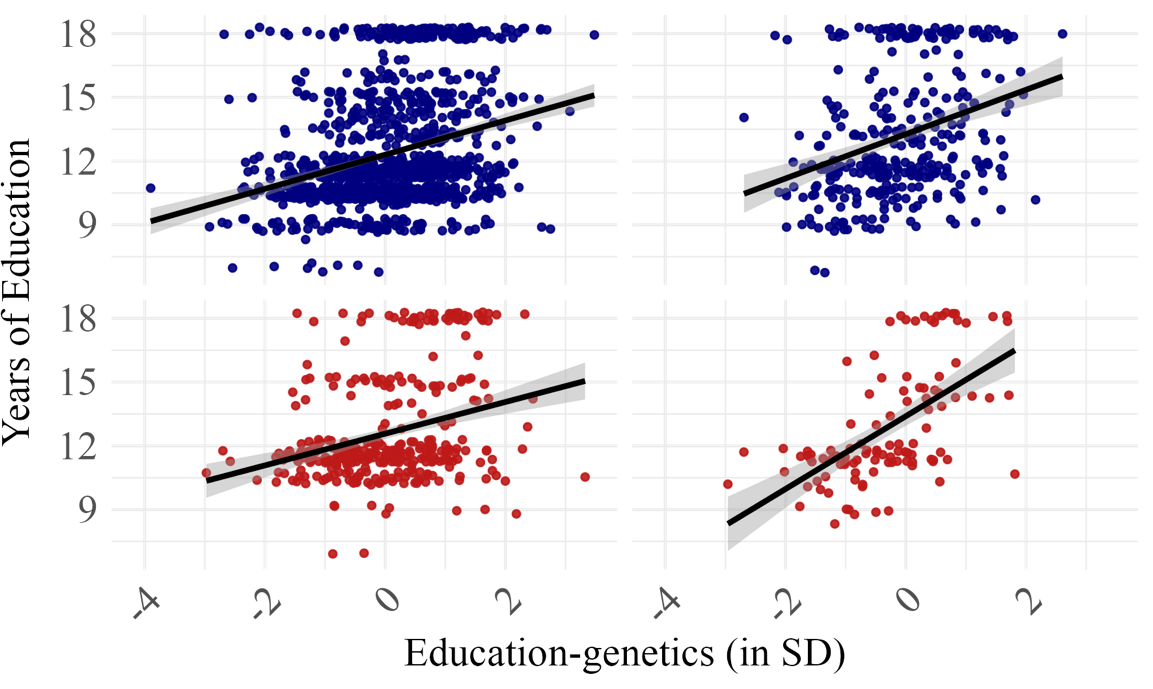

**Figure S2. Scatterplot of the association between education-genetics and educational attainment by region before and after German reunification in East and West Germany.**

| **(A)** | | |
| --- | --- | --- |
|  | 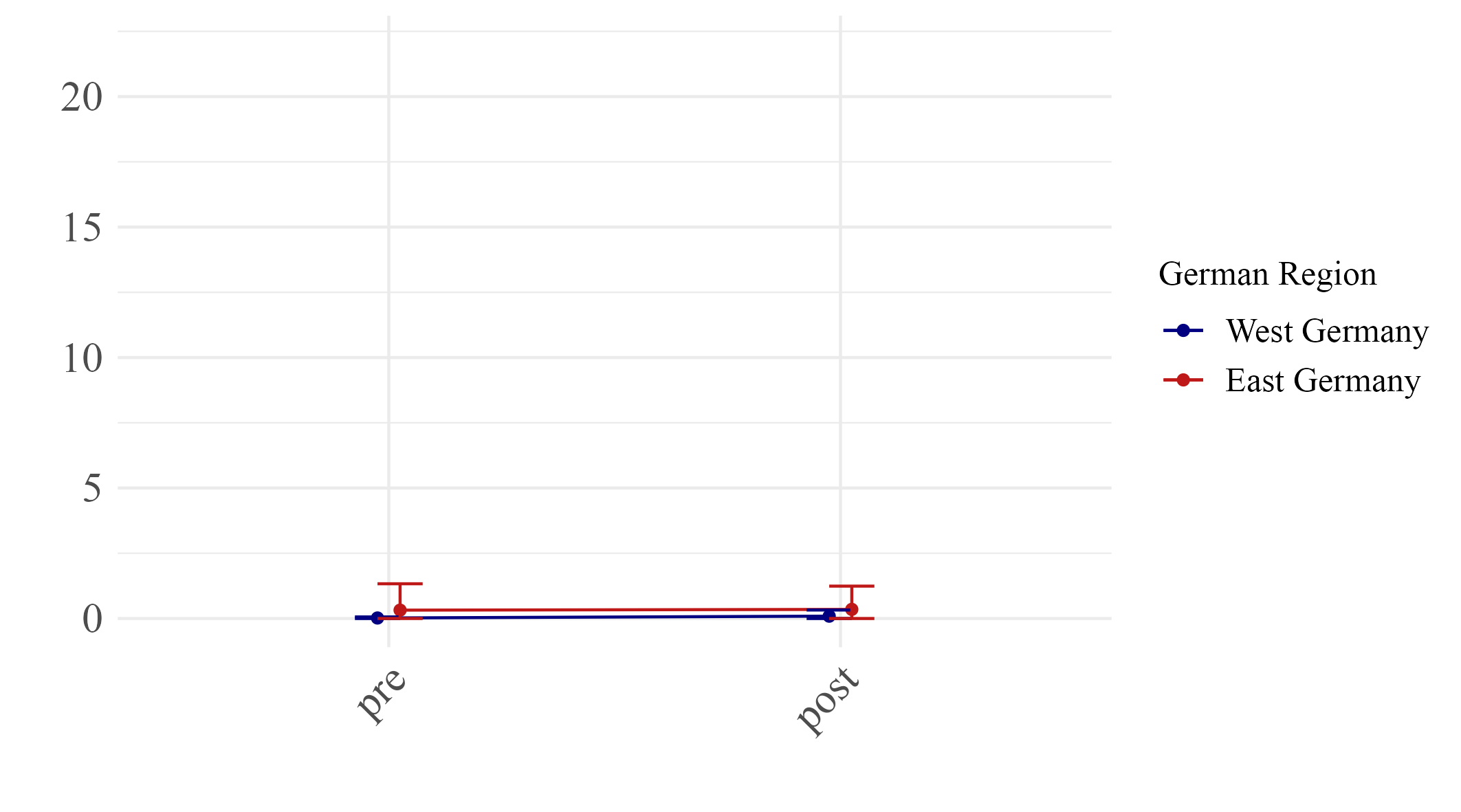 | 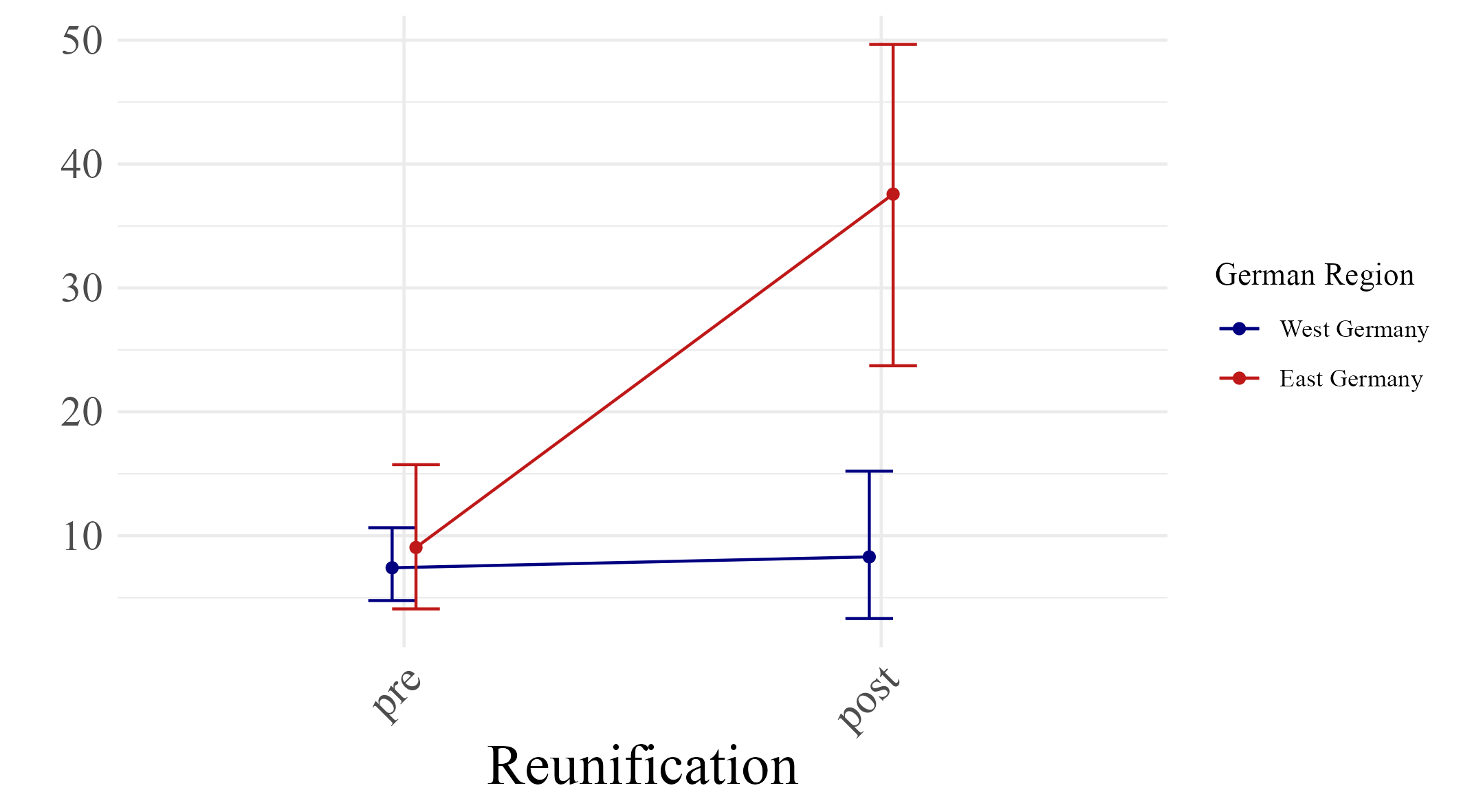 |
|  | Reunification |  |
| **(B)** | 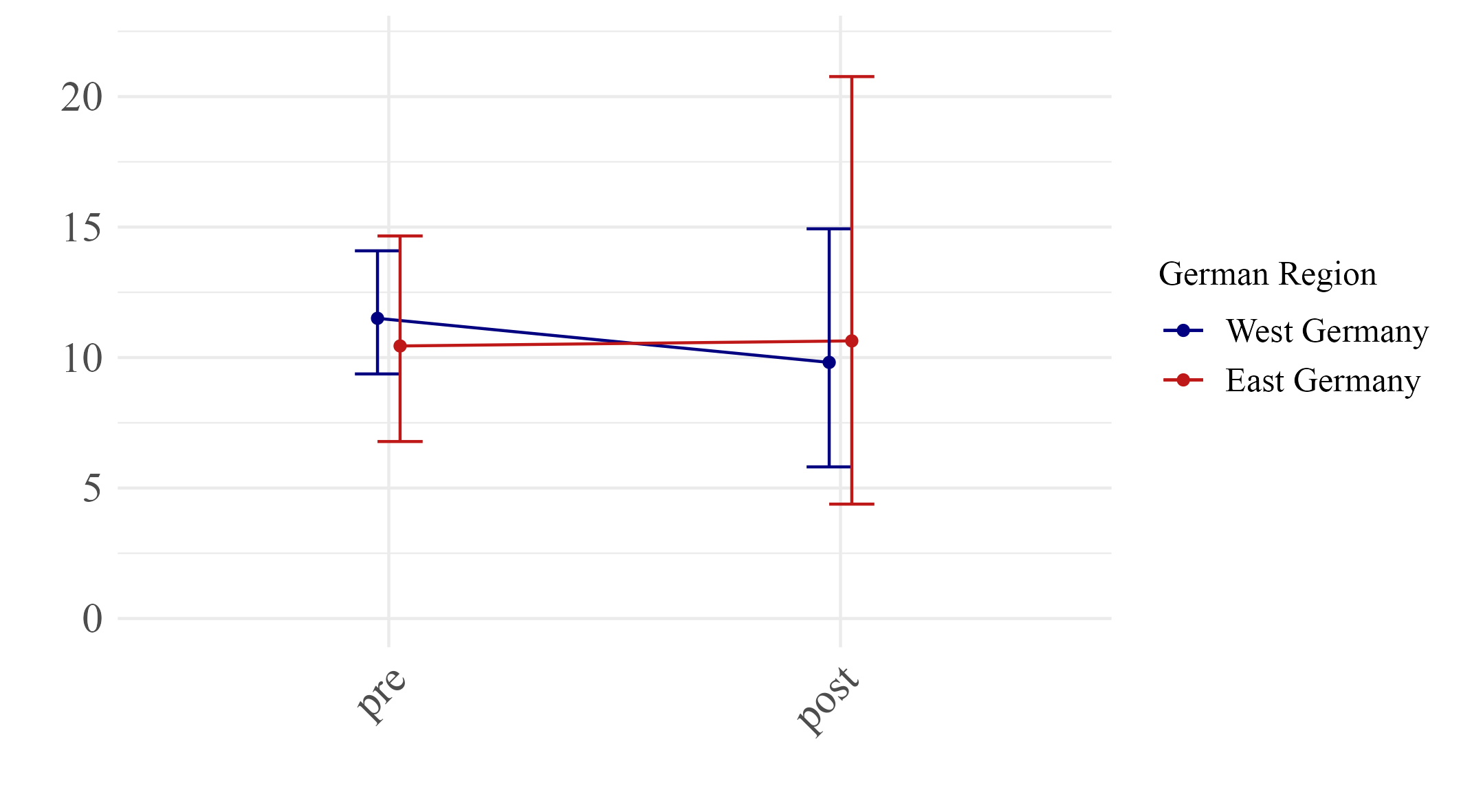  Height  incremental R² (%) |  |
|  | Reunification |  |
| **Figure S3**. Effect size estimates of associations between the polygenic index of height with educational attainment (**Panel A**) and height (**Panel** **B**) before and after German reunification in East and West Germany. | | |

Educational attainment incremental R² (%)
